# Assessment of the *in vitro* function of human stem cell-derived β cells

**DOI:** 10.1101/656785

**Authors:** Arvind R. Srivatsava, Stefanie T. Shahan, Lisa C. Gutgesell, Leonardo Velazco-Cruz, Jeffrey R. Millman

## Abstract

Insulin-producing human embryonic stem cell-derived β (SC-β) cells are a promising cell source for diabetes cell replacement therapy. We have recently reported a differentiation strategy that produces SC-β cells in islet organoids that not only undergo glucose-stimulated insulin secretion but also have an islet-like dynamic insulin release profile, displaying both first and second phase insulin secretion. The goal of this study was to further characterize the functional profile of these SC-β cells *in vitro*. We utilized a Seahorse extracellular flux analyzer to measure mitochondrial respiration of SC-β cells at low and high glucose. We also used photolithography to fabricate a microfluidic device containing microwells to immobilize SC-β cells for perfusional analysis, monitoring cytoplasmic calcium using Fluo-4 AM at low and high glucose. Here we find that in addition to increased insulin secretion, SC-β cells have increased cellular respiration and cytoplasmic calcium ion concentration in response to a high glucose stimulation. Our results indicate that SC-β cells have similar function to that reported for islets, providing further performance characterization that could help with eventual development for diabetes cell therapy and drug screening.

---

Diabetes Mellitus (DM) is a group of metabolic disorders that leads to the inability of the body to regulate blood glucose levels. Type 1 diabetes (T1D) involves the autoimmune-mediated destruction of the insulin-producing pancreatic β cells located in the islets of Langerhans, leading to insulin deficiency and hyperglycemia. T1D is typically managed by injection of exogenous insulin. However, this clinical intervention does not emulate the normal behavior of the native β cells, making patients at risk for many long term complications^1^. An alternative therapy for T1D is the transplantation of cadaveric human islets, with the hope of reducing hyper-and hypoglycemic episodes. Some patients who were transplanted with islets have remained insulin independent for several years^2^. A major limitation of this approach, however, is the scarcity and quality of islets sourced from cadavers, limiting the widespread application of this therapy. Differentiation of human embryonic stem cells (hESCs) to insulin-producing β (SC-β) cells in islet organoids could serve as an unlimited supply of cells to treat millions of patients^3^, particularly if combined with transplantation strategies that vascularization, allow retrievability, and/or protect the cells from immune attack^4-8^.

We recently reported a strategy for making large numbers of SC-β cells from hESCs^9^. This protocol is highly efficient, generating an almost pure population of pancreatic endocrine, of which most cells express insulin. These cells are highly functional, capable of undergoing glucose-stimulated insulin secretion. These cells were capable of restoring glucose tolerance when transplanted into streptozotocin-treated mice. Of particular note was the ability of these cells to display first and second phase insulin secretion kinetics *in vitro* in response to high glucose in a perifusion assay, a feature missing in other strategies.

In β cells found within islets, a high glucose stimulation creates a cascade of intracellular events ultimately leading to the insulin secretion^10^. Mitochondrial respiration is increased, leading to an increase in ATP/ADP rations, a closing of potassium channels, and depolarization of the plasma membrane. This in turn triggers the opening of voltage-gated calcium ion channels, leading to the influx of calcium ions into the cell, inducing the exocytosis of the insulin granules.

The purpose of this study was to further characterize the function of SC-β cells generated with the protocol published in Velazco-Cruz et al^9^. We performed static and dynamic glucose-stimulated insulin secretion to confirm the *in vitro* function of these cells. We observed that the cells were robustly function, capable of undergoing first and second phase insulin secretion in response to a 20 mM glucose stimulation. We then assessed mitochondrial respiration by measuring the oxygen consumption rate using a Seahorse XF24 at low and high glucose and found that the cells increase their respiration in response to 20 mM glucose. Finally, we built a microfluidic device that allows for live fluorescent imaging of our SC-β cells as we perfuse low and high glucose and found cytoplasmic calcium to increase at 20 mM glucose.

## Results

### Generation of SC-β cells for assessment

The goal for this study was to characterize the *in vitro* function of SC-β cells generated with our recent differentiation protocol^9^. This protocol uses six stages with defined, time-specific combinations of small molecule and protein factors to recapitulate pancreatic development to generate functional SC-β cells (**Fig. 1a**). The protocol is performed entirely in suspension culture, producing clusters of endocrine cells that are islet-like in size (**Fig. 1b**). These clusters stain red when treated with the zinc-chelating agent dithizone, a dye known to stain β cells (**Fig. 1c**).

**Figure 1.**
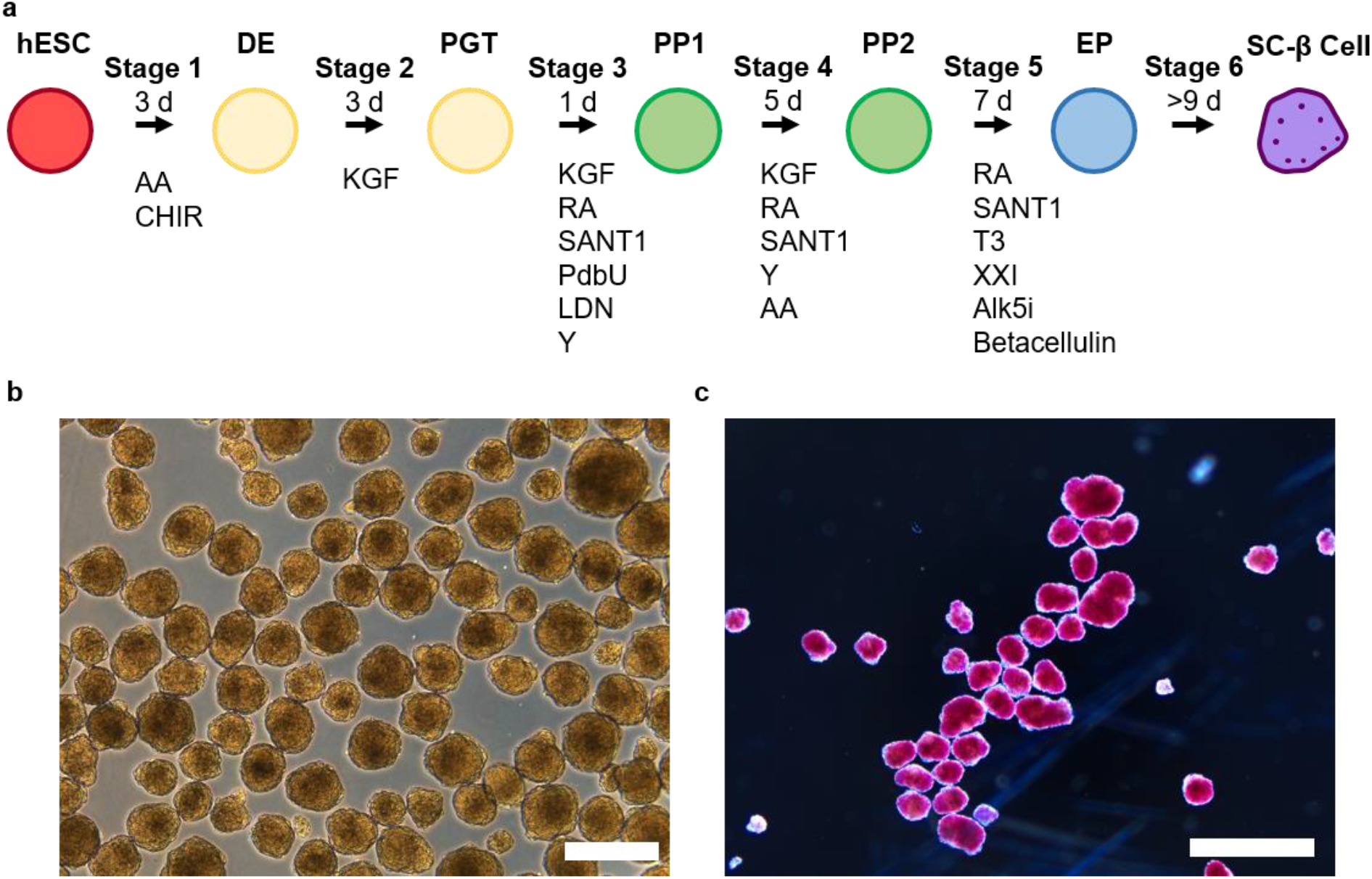
Differentiation of hESCs to SC-β cells. **(a)** Schematic overview of the six-stage protocol. **(b)** Image of stage 6 clusters containing SC-β cells. Scale bar=100 μm. **(c)** Image of dithizone staining of stage 6 clusters, which identifies β cells by staining red when viewed under bright field. Scale bar = 1 mm. DE, definitive endoderm; PGT, primitive gut tube; PP1, pancreatic progenitor 1; PP2, pancreatic progenitor 2; EP, endocrine progenitor; AA, activin A; CHIR, CHIR9901; KGF, keratinocyte growth factor; RA, retinoic acid; Y, Y27632; LDN, LDN193189; PdbU, phorbol 12,13-dibutyrate; T3, triiodothyronine; Alk5i, Alk5 inhibitor type II; ESFM, enriched serum-free medium.

We evaluated the SC-β cells produced with this protocol for expression of β cell markers and capability of undergoing glucose-stimulated insulin secretion. When examined histologically, these clusters in stage 6 express C-peptide, a peptide produce by the insulin gene, and many C-peptide+ cells co-stain with PDX1, a β cell transcription factor (**Fig. 2a**). We assessed the function of the cells with both static and dynamic assays. To perform the static assay, we incubated stage 6 clusters for 1 hr at low (2 mM) glucose, collected the supernatant, then incubated again for 1 hr at high (20 mM) glucose, and collected the supernatant. We observed that these cells secreted 3.0±0.4x more insulin at high compared to low glucose (**Fig. 2b**). To perform the dynamic assay, we perifused solutions of low and high glucose through chambers loaded with the stage 6 clusters, collecting the flow-thru every 2 min. We observed the cells responded to a high glucose challenge with both a first and second phase insulin release profile, with the first phase response occurring within 3-5 min (**Fig. 2c**). These data support that our differentiation protocol produces cells that express β cell markers and are capable of undergoing robust glucose-stimulated insulin secretion.

**Figure 2.**
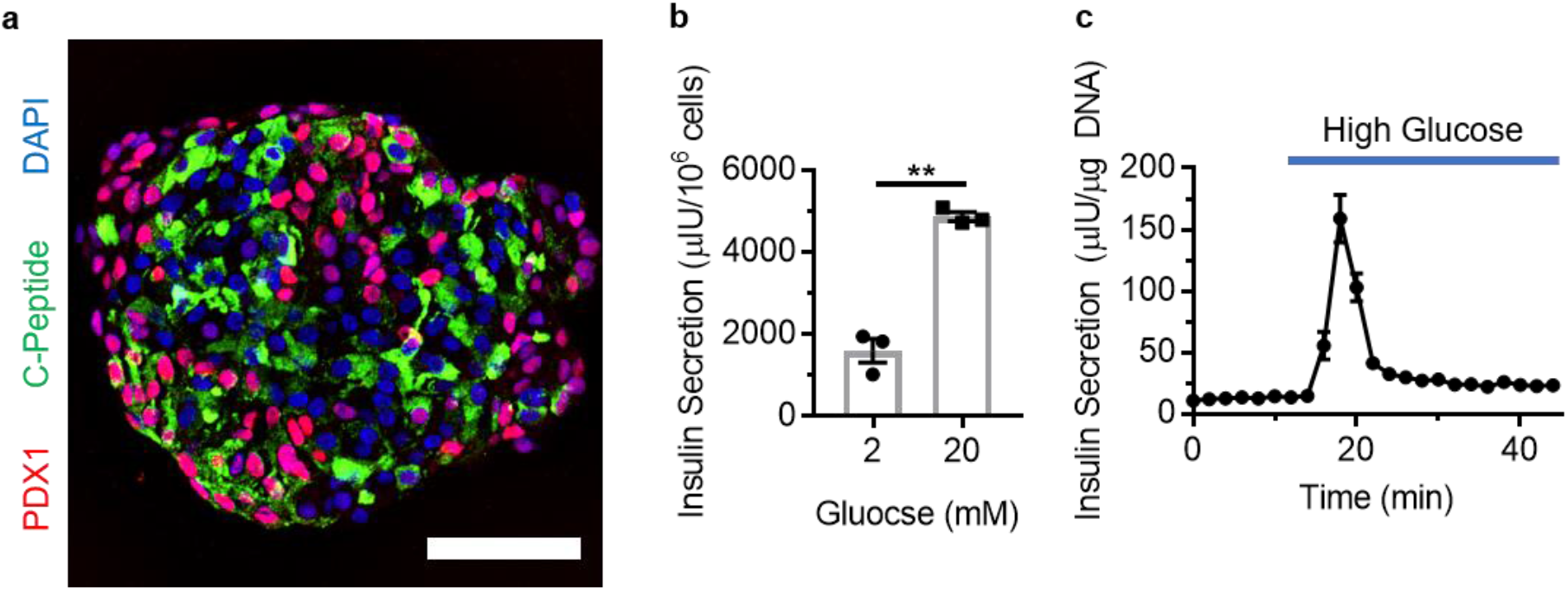
SC-β cells express β cell markers and undergo glucose-stimulated insulin secretion. **(a)** Immunostaining of sectioned stage 6 cluster stained for C-peptide, which is produced by the insulin gene, and PDX1, a β cell transcription factor. **(b)** Static glucose-stimulated insulin secretion. **p<0.01 by paired two-way **t**-test. n=3. **(c)** Dynamic glucose-stimulated insulin secretion. n=3.

### Mitochondrial respiration of SC-β cells

In order to better characterize SC-β cells produced with our protocol, we assessed their mitochondrial respiration. As part of the molecular machinery for sensing glucose, β cells have increased mitochondria respiration with increasing glucose^11^. To assess our cells, we used an Seahorse XF24 instrument^12^, which is capable of simultaneously measuring oxygen consumption rate (OCR), used to monitor mitochondria respiration, and extracellular acidification rate (ECAR), used to monitor glycolysis. We loaded 20-30 hESC or SC-β cell clusters into low glucose-filled microwells compatible with the instrument (**Fig. 3a**). After establishing a baseline for respiration, we injected high glucose and observed a 56±5% increase in OCR for SC-β cells, with only a minor response observed with hESCs (**Fig. 3b**). Subsequent injections were made with compounds to further interrogate mitochondria respiration (**Fig. 3b)**: Oligomycin (OM), which binds and inhibits ATP synthase, Carbonyl cyanide-*4*-(trifluoromethoxy)phenylhydrazone (FCCP), which is an uncoupler, transporting protons into the mitochondria, and Antimycin A (AA) and rotenone (R), which inhibit cellular respiration. We observed that the SC-β cells had a much greater reduction in OCR with OM treatment due to their elevated respiration at high glucose and a greater response to FCCP treatment, both of which indicate greater respiratory capacity of these cells compared to hESCs. OCR was also greatly reduced with AA/R treatment for both cell types, indicating that non-mitochondrial sources of respiration make up only a small portion of OCR. These data show that SC-β cells respond to high glucose by increasing mitochondrial respiration.

**Figure 3.**
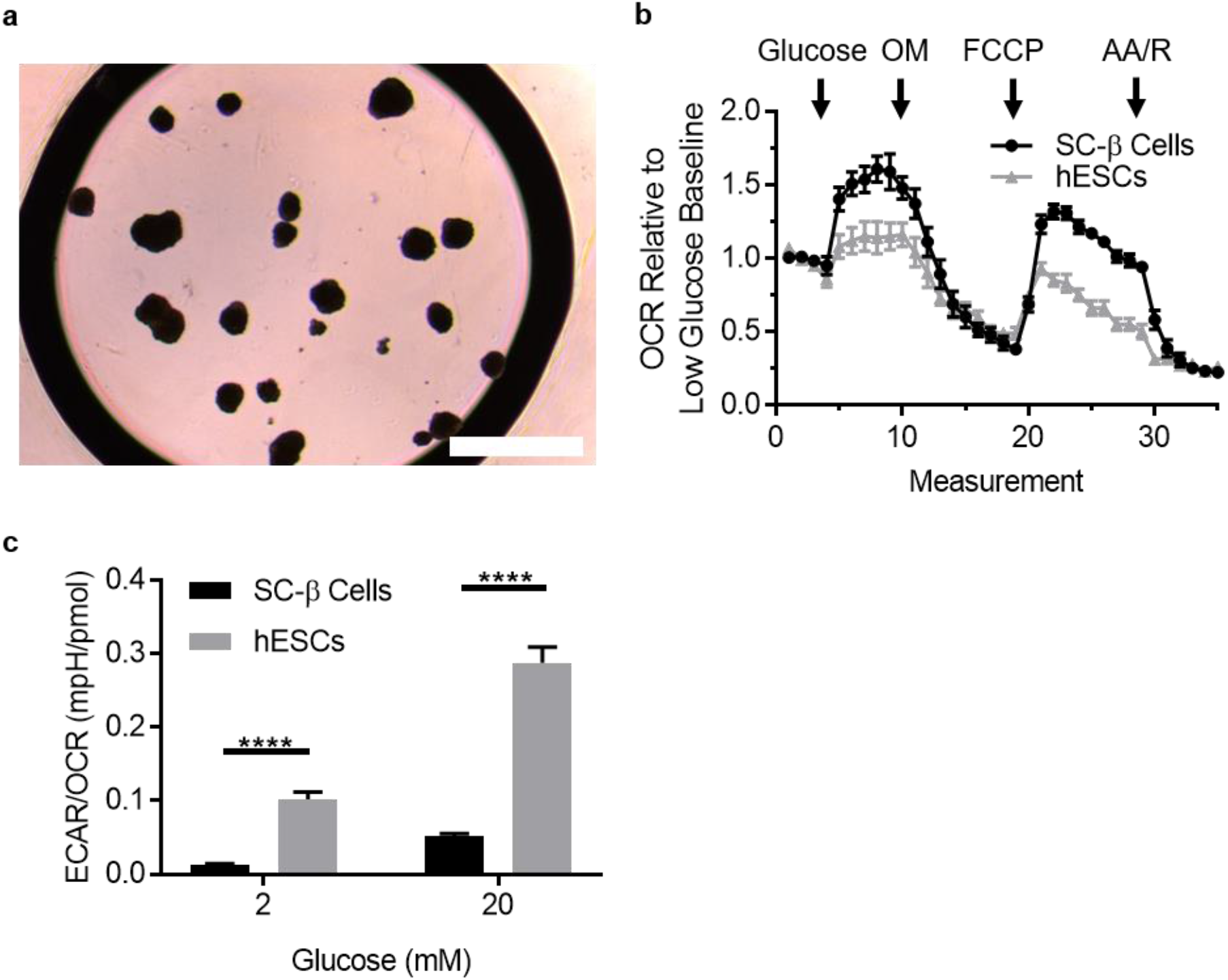
SC-β cells have elevated mitochondrial respiration. **(a)** Image of clusters placed into microwell for Seahorse XF24 analysis. Scale bar = 1 mm. **(b)** Measurements of oxygen consumption rate (OCR) for 35 sequential measurements with injection of indicated compounds. **(c)** Ratio of extracellular acidification rate (ECAR) to OCR at low and high glucose.

To further interrogate SC-β cell metabolism, we assessed the ratio of glycolysis to mitochondrial respiration, as defined by the ECAR/OCR ratio. Measuring these ratios at low and high glucose for SC-β cells and hESCs, we observed hESCs to have a much higher ratio, indicating relatively higher glycolysis in hESCs (**Fig. 3c**). In addition, we used real-time PCR to measure the gene expression differences of SC-β cells and hESCs and observed gene expression of mitochondrial transporters related to oxidative phosphorylation were higher in SC-β cells, correlating with increased INS and lower OCT4 transcript expression (**Fig. 4**). In contrast, LDHA gene expression, which is necessary to sustain high levels of glycolysis, decreased with differentiation to SC-β cells (**Fig. 4**). Taken together, these data show that differentiation of hESCs to SC-β cells increases respiratory capacity along with expression of many associated genes and decreases glycolysis along with LDHA gene expression.

**Figure 4.**
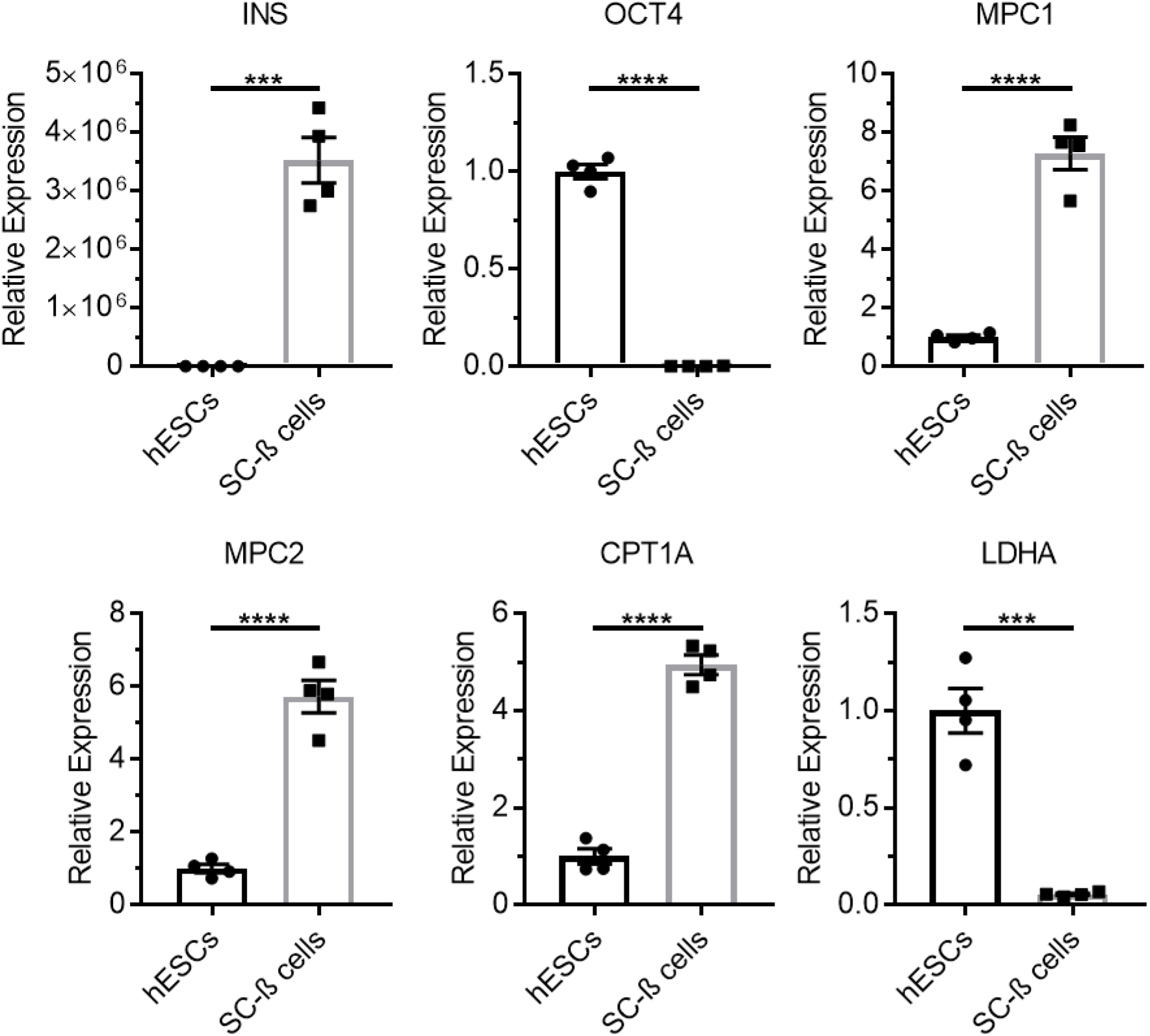
Gene expression of metabolic, β cell, and hESCs genes. Relative gene expression was measured with real-time PCR. ***p<0.001, ****p<0.0001. n=4.

### Fabrication of microfluidic device for assessing cytoplasmic calcium

To facilitate study of cytoplasmic calcium levels in SC-β cells, we fabricated a microfluidic device^13^. The microfluidic device for this experiment is comprised of three layers (**Fig. 5a**): A bottom layer containing microwells 500 μm in diameter and 150 μm in depth to immobilize cellular clusters, a middle layer containing the microchannel measuring 19 mm x 2 mm x 250 μm and a reservoir cutout 7 mm in diameter and 5 mm in depth, and a top transparent layer that covers the first two layers. We drew these designs using AutoCAD to create photomasks for the lithography process using a Laser Writer (**Fig. 5b**). These masks were used to make a master with SU-8 photoresist (**Fig. 5c**). We then proceeded with soft lithography by spin coating the master with Polydimethylsiloxane (PDMS) that is then cured and each layer assembled (**Fig. 5d**). In the end, this produced a microfluidic device with microwells for immobilizing cellular clusters (**Fig. 5e**).

**Figure 5.**
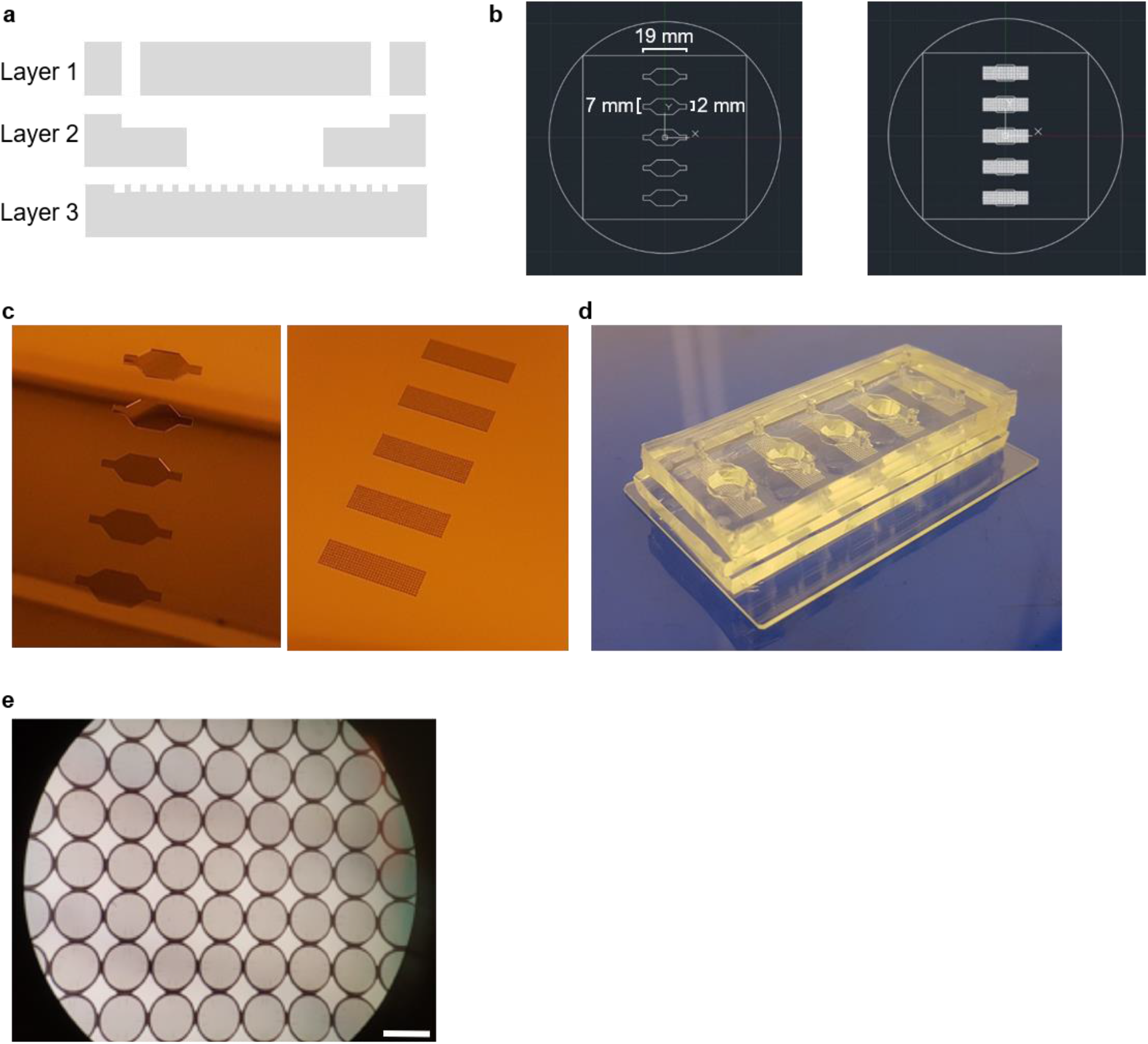
Manufacture and assembly of microfluidic device for cytoplasmic calcium ion imaging. **(a)** Illustration of the three-layer device, with the top layer containing the inlet and outlet, the middle layer containing the channel and fluid reservoir, and the bottom layer containing the microwells to immobilize cellular aggregates. **(b)** AutoCAD drawing of layers 2 and 3 used to make the photomasks. **(c)** Images of the fabricated masters for layers 2 and 3. **(d)** Image of the finished and assembled microfluidic device made out of PDMS. **(e)** Micrograph of the microwells in the bottom of the microfluidic device. Scale bar = 500 μm.

Next, we tested the microfluidic device with a setup consisting of syringe pumps to continuously flow buffers at 0.5 mL/min with defined glucose concentrations into the device and a fluorescence microscope to image cells stained with Fluo-4 AM, a dye that fluorescences proportional to cytoplasmic calcium concentration (**Fig. 6a**). We first tested the glucose switching action of the experimental setup, switching from low to high glucose and sampling the flow through every two min. We observed a rapid step change in the outline concentration of glucose, establishing the fast glucose switching action of the setup that will enable us to proceed further with using cells in the device (**Fig. 6b**). We next loaded cells into the device and confirmed that the clusters were immobilized within the microwells as designed in layer 3 (**Fig. 6c**). Finally, we stained SC-β cells with Fluo-4 AM and were able to successfully image the cells within the device (**Fig. 6d**). These data provided validation of the experimental setup that enabled experimentation studying cytoplasmic calcium in SC-β cells.

**Figure 6.**
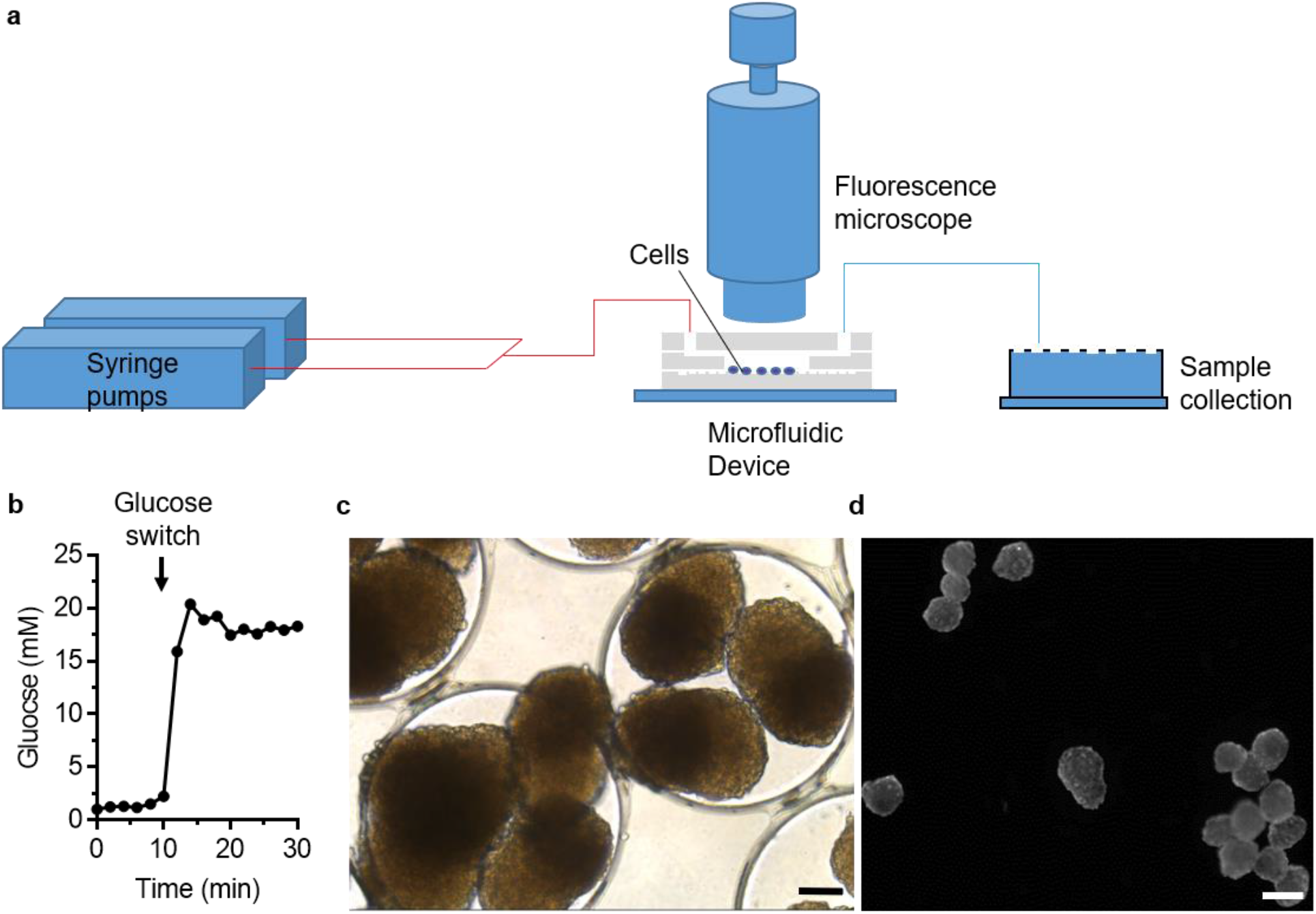
Loading of cells into the microfluidic device. **(a)** An illustration of the microfluidic perfusion setup that allows for imaging of Fluo-4 AM-stained cell clusters. **(b)** Perfusion of glucose switched from 2 to 20 mM. **(c)** Bright field image of stage 6 clusters immobilized in the micowells within the device. Scale bar = 100 μm. **(d)** Fluorescence image of Fluo-4 AM-stained stage 6 clusters loaded into the microfluidic device. Scale bar = 200 μm.

### Cytoplasmic calcium of SC-β cells

To further our characterization of SC-β cells produced with our protocol, we assessed their cytoplasmic calcium levels. As part of the molecular machinery for sensing glucose, β cells have increased cytoplasmic calcium with increasing glucose^14^. We perfused SC-β cell clusters under low glucose, high glucose, and high KCl, continuously monitoring Fluo-4 AM fluorescence. We monitored 16 individual clusters that were within the view of the microscopy and plotted the responses of each cluster in **Fig. 7a**. We observed that all 16 SC-β cell clusters to respond to the switch to high glucose by increasing Fluo-4 AM fluorescence by an average factor of 1.120±0.002. This increase was even greater with the addition of KCl, which depolarizes the membrane and causing calcium ion influx into β cells, increasing Fluo-4 AM fluorescence by an average factor of 1.248±0.010 compared to low glucose. We collected the flow-thru every 3 min and quantify secreted insulin concentration, plotting these concentrations to the average Flou-4 AM fluorescence in **Fig. 7b**. In addition to increases in Fluo-4 AM fluorescence with high glucose and KCl treatment, insulin secretion was also increased with these stimulations. KCl treatment had larger increases in both insulin and cytoplasmic calcium than high glucose. These data show that SC-β cells respond to high glucose by increasing cytoplasmic calcium, which correlates with insulin secretion.

**Figure 7.**
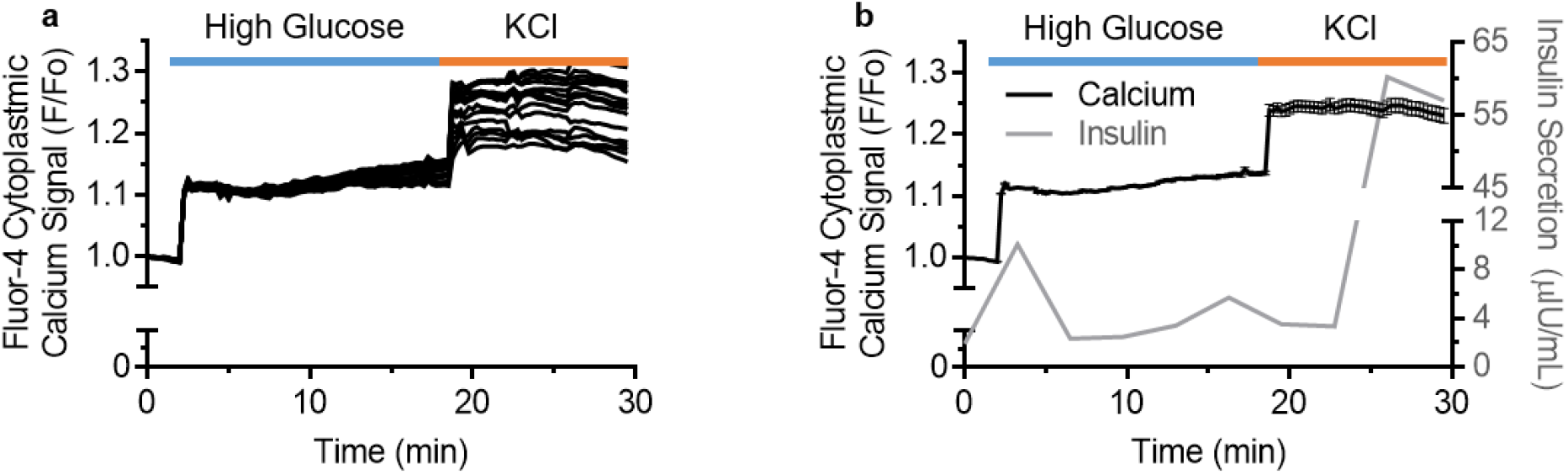
Quantification of cytoplasmic calcium changes in SC-β cell clusters. **(a)** Traces of 16 individual SC-β cell clusters measuring the change in Fluo-4 AM fluorescence intensity going from low (2 mM) to high (20 mM) glucose and to high KCl. **(b)** Averaged change in Fluo-4 AM fluorescence intensity plotted against insulin concentration of the collected flow-thru.

## Discussion

In this study, we evaluated the functional characteristics of SC-β cells generated with our 6-stage differentiation protocol (**Fig. 8**). We found that these cells, in addition to expressing markers of β cells, undergo glucose-stimulated insulin secretion and have robust dynamic function, displaying first and second phase insulin secretion kinetics to a high glucose stimulation. This insulin secretion is associated with increases in mitochondrial respiration, which were higher than undifferentiated hESCs. Genes associated with mitochondrial respiration but not glycolysis were higher in SC-β cells than hESCs. In addition, this function was associated with increases in cytoplasmic calcium levels, which we assessed by fabricating and validating a microfluidic device that perfuses Fluo-4 AM-stained SC-β cell clusters with buffers of defined glucose for real-time imaging with a fluorescence microscope.

**Figure 8.**
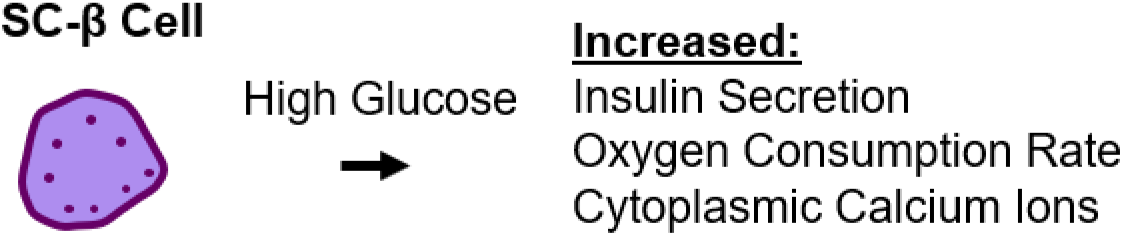
Schematic overview of major findings from the study.

Proper mitochondrial function within β cells is key to their health and function. The process of insulin release relies on the mitochondria to process pyruvate and produce ATP. Once ATP has been produced, it inhibits the outward flow of potassium by blocking potassium channels and depolarizes the cell, causing an inward flow of calcium that then stimulates insulin granule exocytosis. Without proper functioning of the mitochondria, insulin release can be disrupted. The disruption of insulin release due to an error in this process can lead to diabetes. Notably, mutations in mitochondrial DNA (mitochondria have their own set of DNA, separate from the nucleus of the cell they reside in) can initiate diabetes, with no other known mutation^15^, implying that a disruption in solely mitochondrial function is enough to cause diabetes and highlighting the importance of healthy mitochondria to β-cell function.

One distinct characteristic of hESCs that differs from mature cells is their metabolism: they are heavily reliant on glycolysis, an anaerobic process that generates ATP using glucose, to generate their energy^16^. Accordingly, their mitochondria differ from other cell types in that hESCs possess scarce mitochondria that are localized to the edges of the cell, are underdeveloped with poor cristae formation, and appear to have restricted oxidative capacity^17^. As hESCs differentiate, their mitochondrial networks increase in mass, branch out, and localize to other areas of the cell^18^, and their mitochondria become more active as oxidative phosphorylation becomes the primary source of energy^19^. Despite induced pluripotent stem cells (iPSCs) being generated from adult cells with developed mitochondria and a reliance on oxidative phosphorylation, they seem to revert to a state similar to hESCs in terms of metabolism and mitochondria^20^ and, upon differentiation, share similar mitochondrial regulation mechanisms as hESCs^17,21^. Altering hESC mitochondrial biogenesis has been shown to influence commitment to a germ layer in differentiation^22^. Calcium is known to help regulate mitochondrial respiration in β cells^23^.

The SC-β cells work similarly to β cells from human islets in sensing changing glucose concentration^9^. Insulin secretion response is typically characterized to have two phases^24^, the first phase being a large spike occurring within minutes after glucose stimulation, attributed to the exocytosis of readily releasable insulin granules close to the plasma membrane which release insulin in response to nutrient and non-nutrient secretagogues^25^, followed by the second phase of insulin release, where the insulin quantity reduces to a suprabasal level and continues to increase gradually as long as external glucose stimulation persists. The second phase of secretion is explained by the intracellular granules or vesicles mobilizing and beginning to dock and fuse with the t-SNARE sites on the plasma membrane^25^, from where insulin exits the cells.

Since glucose-stimulated insulin secretion is directly linked to calcium ion flux, direct analysis of this behavior is desirable. We utilized a microfluidic approach to assess calcium in our SC-β cells^13^. Microfluidics can be defined as the science of study of fluids through micro-channels and as the technology of manufacturing micro-miniaturized devices containing various chambers and convolutions through which fluids are confined^26^ and has been used to generate a wide array of particles^27^. Static analysis of islet function can be achieved through islet multiplexing in a multi-well plate, although this being a static method only enables measurement of bulk insulin produced by the cells (secretory capacity) and does not provide insight on the temporal dynamics of insulin secretion, comprising of the biphasic response of insulin secretion over time that can be achieved with a perfusion microfluidic assessment. The microfluidic approach is also compatible with fluorescence imaging, which is a non-invasive method for real-time data acquisition. We leveraged this capability to image cytoplasmic calcium with Fluo-4 staining. Fluo-4 finds use in many applications such as flow cytometry, including microscopy and microplate screening assays^28^. With regard to this study, it serves as an excellent assessment tool owing to its high rates of cell permeation and the ability to quantify cellular Ca^2+^ changes over a very broad range of concentration.

Major translational applications of SC-β cells is for diabetes cell therapy and drug screening^29^. Over 20 million people have diabetes in the US, and a current need in the field is for translation of stem cell technology for cell therapy is the optimization of differentiation and functional characterization of the resulting cells. Differentiation can be costly, plagued with cell death and lack of efficiency, and take extensive culture time to complete. Optimizing the process to reduce these issues is essential for the transition of stem cells from research subject to plausible, available medical therapy. Differentiation is often altered by changing factors and their concentrations, the duration spent in certain differentiation steps, alteration of cell cluster sizes, or oxygen availability; however, even the best modern differentiations have much room for improvement. With the current progress of research in this field, SC-β cells could be used a cell source in the near future, which meet the current need for cell sourcing, and a multi-faceted approach to analyzing the function of these cells that includes glucose-stimulated insulin secretion, mitochondrial respiration, and cytoplasmic calcium measurements with a microfluidic approach could be key to properly characterizing these cells. This would synergize with other efforts in the field to study SC-β cells or earlier progenitors for disease modeling purposes, particularly with patient-derived cells^30-42^.

## Conclusions

Out data shows that SC-β cells generated with our protocol exhibits strong glucose-stimulated insulin secretion. This function correlates with increases in islet-like mitochondrial respiration and cytoplasmic calcium levels in response to high glucose stimulation. In addition, expression of genes associated with mitochondria respiration increased while glycolysis decreased with differentiation. These findings indicate that SC-β cells are glucose-responsive by using a similar mechanism of β cells found in islets and provide important additional characterization information of their functional potential for diabetes cell replacement therapy and drug screening.

## Acknowledgements

This work was supported by the NIH (R01DK114233), JDRF Career Development Award (5-CDA-2017-391-A-N), Washington University Center of Regenerative Medicine, and startup funds from Washington University School of Medicine Department of Medicine. L.C.G. was supported by the Amgen Scholars Program. L.V.C. was supported by the NIH (R25GM103757). Microscopy was performed through the Washington University Center for Cellular Imaging (WUCCI), which is supported by Washington University School of Medicine, CDI (CDI-CORE-2015-505), and the Foundation for Barnes-Jewish Hospital (3770). The Washington University Diabetes Research Center (P30DK020579) provided support for use of the Seahorse XF24.

The Washington University in St. Louis and the Institute of Materials Science and Engineering provided support for the use of instruments and staff assistance in the microfluidic device fabrication and particularly thank Dr. Rahul Gupta for assistance.

## Author Contributions

A.R.S., S.T.S., and J.R.M. conceived of the experimental design. All authors contributed to the experiments. A.R.S., S.T.S., and J.R.M. wrote the manuscript. All authors edited and reviewed the manuscript.

## Methods and Materials

### Stem cell culture and differentiation to SC-β cells

All work was performed with the HUES8 hESC line, generously provided by Dr. Douglas Melton (Harvard University) and has been previously published on ^3,9^. All hESC culture and differentiation was performed in 100-mL or 30-mL spinner flasks (REPROCELL; ABBWVS10A or ABBWVS03A) on a stirrer plate (Chemglass) set at 60 RPM in a humidified incubator set at 5% CO_2_ and 37 °C. Undifferentiated hESCs were cultured in mTeSR1 (StemCell Technologies; 05850) in Accutase (StemCell Technologies; 07920) and passaged every 3 d.

hESCs were differentiated to SC-β cells as previously described^9^ by culturing in the following conditions in order: <u>Stage 1</u>: 3 d with 100 ng/ml Activin A (R&D Systems; 338-AC) and 3 μM CHIR99021 (Stemgent; 04-0004-10) for 1 d followed with 100 ng/ml Activin A for 2 d. <u>Stage 2</u>: 3 d with 50 ng/ml KGF (Peprotech; AF-100-19). <u>Stage 3</u>: 1 d with 50 ng/ml KGF, 200 nM LDN193189 (Reprocell; 040074), 500 nM PdBU (MilliporeSigma; 524390), 2 μM Retinoic Acid (MilliporeSigma; R2625), 0.25 μM Sant1 (MilliporeSigma; S4572), and 10 μM Y27632. <u>Stage 4</u>: 5 d with 5 ng/mL Activin A, 50 ng/mL KGF, 0.1 μM Retinoic Acid, 0.25 μM SANT1, and 10 μM Y27632. <u>Stage 5</u>: 7 d with 10 μM ALK5i II (Enzo Life Sciences; ALX-270-445-M005), 20 ng/mL Betacellulin (R&D Systems; 261-CE-050), 0.1 μM Retinoic Acid, 0.25 μM SANT1, 1 μM T3 (Biosciences; 64245), and 1 μM XXI (MilliporeSigma; 595790). <u>Stage 6</u>: >9 d without factors. <u>Stage 1 base media was</u>: 500 mL MCDB 131 (Cellgro; 15-100-CV) plus 0.22 g glucose (MilliporeSigma; G7528), 1.23 g sodium bicarbonate (MilliporeSigma; S3817), 10 g bovine serum albumin (BSA) (Proliant; 68700), 10 μL ITS-X (Invitrogen; 51500056), 5 mL GlutaMAX (Invitrogen; 35050079), 22 mg vitamin C (MilliporeSigma; A4544), and 5 mL penicillin/streptomycin (P/S) solution (Cellgro; 30-002-CI). <u>Stage 2 base media was</u>: 500 mL MCDB 131 plus 0.22 g glucose, 0.615 g sodium bicarbonate, 10 g BSA, 10 μL ITS-X, 5 mL GlutaMAX, 22 mg vitamin C, and 5 mL P/S. <u>Stage 3 and 4 base media was</u>: 500 mL MCDB 131 plus 0.22 g glucose, 0.615 g sodium bicarbonate, 10 g BSA, 2.5 mL ITS-X, 5 mL GlutaMAX, 22 mg vitamin C, and 5 mL P/S. <u>Stage 5 base media was</u>: 500 mL MCDB 131 plus 1.8 g glucose, 0.877 g sodium bicarbonate, 10 g BSA, 2.5 mL ITS-X, 5 mL GlutaMAX, 22 mg vitamin C, 5 mL P/S, and 5 mg heparin (MilliporeSigma; A4544). <u>Stage 6 base media was</u>: 500 mL MCDB 131 plus 0.23 g glucose, 10.5 g BSA, 5.2 mL GlutaMAX, 5.2 mL P/S, 5 mg heparin, 5.2 mL MEM nonessential amino acids (Corning; 20-025-CI), 84 μg ZnSO_4_ (MilliporeSigma; 10883), 523 μL Trace Elements A (Corning; 25-021-CI), and 523 μL Trace Elements B (Corning; 25-022-CI). Clusters at the end of stage 5 were reaggregated by dispersion with TrypLE Express (ThermoFisher; 12604013).

### Light microscopy

Bright field images of cells were taken with an inverted light microscope (Leica DMi1). DTZ staining was done at 2.5 μg/mL and purchased from MilliporeSigma (194832).

### Immunostaining

Whole clusters were fixed with 4% paraformaldehyde (Electron Microscopy Science; 15714), embedded in Histogel to aid with sectioning (Thermo Scientific; hg-4000-012), embedded in paraffin and sectioned by the Division of Comparative Medicine (DCM) Research Animal Diagnostic Laboratory Core at Washington University. To immunostain, paraffin was removed with Histoclear (Thermo Scientific; C78-2-G), the samples rehydrated, and samples treated with 0.05 M EDTA (Ambion; AM9261) in a pressure cooker (Proteogenix; 2100 Retriever) to retrieve antigens. Blocking was performed with a 30-min treatment of staining buffer (5% donkey serum (Jackson Immunoresearch; 017-000-121) and 0.1% Triton-X 100 (Acros Organics; 327371000) in PBS). Primary antibodies (1:300 dilutions of rat-anti-C-peptide (DSHB; GN-ID4-S) and goat anti-PDX1 (R&D Systems; AF2419) were left on overnight, after which the samples were stained with secondary antibodies containing Alexa Fluor fluorophores (Invitrogen) for 2 hr and mounted with Fluoromount-G (SouthernBiotech; 0100-20). Imaging was performed on a Nikon A1Rsi confocal microscope.

### Static glucose-stimulated insulin secretion

This assay was performed in KRB buffer consisting of 128 mM NaCl, 5 mM KCl, 2.7 mM CaCl_2_ 1.2 mM MgSO_4_, 1 mM Na_2_HPO_4_, 1.2 mM KH_2_PO_4_, 5 mM NaHCO_3_, 10 mM HEPES (Gibco; 15630-080), and 0.1% BSA. After preincubating the cells at 2 mM glucose, clusters were incubated for 1 hr at 2 mM glucose followed by 1 hr at 20 mM glucose. Insulin in the supernatant from these 1 hr incubations was quantified with a Human Insulin ELISA (ALPCO; 80-INSHU-E10.1). Viable cell numbers were determined with the the Vi-Cell XR (Beckman Coulter).

### Dynamic glucose-stimulated insulin secretion

This assay was performed in KRB buffer as we have previously published^9^ using a perifusion system we assembled based on Bentsi-Barnes et al^43^. Cells were placed in a perifusion chamber (BioRep; Peri-Chamber) and maintained at 37 °C in a water bath. The chamber was constantly perfused with KRB at 100 μl/min. After a pre-incubation at 2 mM glucose, flow-thru was collected every 2 min with switching from 2 to 20 mM glucose. Insulin in the flow-thru was quantified using the Human Insulin Elisa kit, and DNA was quantified using Quant-iT Picogreen dsDNA assay kit (Invitrogen; P7589).

### Mitochondrial respiration

The Seahorse XF24 extracellular flux analyzer (Agilent) were used to measure OCR as we have previously reported^12^. Stage 6 clusters were loaded into islet capture microplates (Agilent; 101122-100) in RPMI with 2 mM glucose. After equilibration and calibration, baseline measurements of OCR and ECAR was taken. Then the following injections were made in sequence: 20 mM glucose, 3 μM oligomycin (Calbiochem; 1404-19-9), 0.25 μM carbonyl cyande-4-(trifluoromethoxy) phenylhydrazone (FCCP) (Sigma; 270-86-5), and 1 μM rotenone (Calbiochem; 83-79-4) and 2 μM antimycin A (Sigma; 1397-94-0). Measurements of OCR and ECAR were taken after each injection.

### Real-time PCR

The RNeasy Mini Kit (Qiagen; 74016) with DNase treatment (Qiagen; 79254) was used to extract RNA from hESCs and Stage 6 cells. cDNA was made with the High Capacity cDNA Reverse Transcriptase Kit (Applied Biosystems; 4368814). Real-time PCR was performed using PowerUp SYBR Green Master Mix (Applied Biosystems; A25741) with a StepOnePlus instrument (Applied Biosystems) and the generated data analyzed using δδCt methodology. TBP was used as the normalization gene. The following primer pairs were used: <u>INS</u>, CAATGCCACGCTTCTGC, TTCTACACACCCAAGACCCG; <u>TBP</u>, GCCATAAGGCATCATTGGAC, AACAACAGCCTGCCACCTTA; <u>LDHA</u>, GGCCTGTGCCATCAGTATCT, GGAGATCCATCATCTCTCCC; <u>OCT4</u>, GGTTCTCGATACTGGTTCGC, GTGGAGGAAGCTGACAACAA; <u>MPC1</u>, TTTTCATATCATTGATGGCAGC, GGACTATGTCCGAAGCAAGG; <u>MPC2</u>, CACCAACCCCCATTTCATAA, TAAAGTGGAGCTGATGCTGC; <u>CPT1A</u>, GCCTCGTATGTGAGGCAAAA, TCATCAAGAAATGTCGCACG.

### Microfluidic device fabrication and assembly

The microfluidic device was based on Mohammed et al^13^. Fabrication was performed at the Institute of Materials Science and Engineering. The photomasks were designed using AutoCAD 2018 software. Designs for both layers were modeled after 50mm diameter sized wafers. Design files were then used to direct the patterns drawn on the photomask substrates by the Laser Writer to generate the photolithographic masks for selective photoresist exposure. For layer 1, photoresist SU8 −2050 was applied on a silicon wafer of diameter 50 mm and spin coated at 1300 RPM for 35 seconds. The coated wafer was then baked on a hot plate for 5 min at 65 °C and then for a longer duration of 30 min at 95 °C. After baking, the coated wafer was placed in a Mask Aligner where the photomask prepared earlier through Laser Writing is mounted in the Aligner. The wafer was then exposed to UV light at an exposure energy of 290 mJ/cm^2^ for 3 min. The mask and wafer were removed, and the exposed wafer was baked (post exposure baking) for 5 min at 65 °C and then for 45 min at 95 °C before being immersed in and treated with SU-8 developer solution with agitation for about 30 min. The developed wafer was then rinsed with isopropanol and water and air dried. For layer 2, the photoresist SU8-2050 was spin coated at 500 RPM for 35 seconds and then soft baked at 65 °C for 7 min, followed by baking at 95 °C for 5 min. The coated wafer was then exposed to UV light at an exposure energy of 385 mJ/cm^2^ before undergoing post exposure baking at 65 °C for 5 min and 95 °C baking for 15 min. The baked wafer was then treated with SU8 developer solution for about 30 min and then rinsed with isopropanol and water. The finished and developed wafer was then air dried.

To fabricate the microfluidic device, 30 g of Sylgard 184 base elastomer was mixed with 3 g of Sylgard 184 elastomer curing agent for a mixing ratio of 10:1 separately in petridishes containing the patterned wafers for Layer 1 and Layer 2 and an empty petridish for Layer 3. The mixture was vigorously stirred until bubbles can be seen. The curing mixture was then deaerated in a vacuum desiccator for long enough until all the bubbles in the mixture disappear. Each of the petridishes are then cured in an oven overnight at 70 °C. After curing, using scalpels, the patterned polymer was shaped and removed from the petri dishes and inlet holes for Layer 3 and the reservoir hole for Layer 2 was punched out using hole-punches. The polymer slabs and a glass slide (to stick the layers) were then immersed in 70% ethanol and sonicated in a water bath for 20 min, to disinfect and remove impurities. As a final step, in order to bond the layers together along with the slide, the slabs were exposed to UV ozone plasma in an Asher/Plasma Cleaner for 30 s at 100W with the surfaces to be bonded being exposed. The layers and the glass slide were then aligned and firmly pressed together to complete the process of device fabrication.

### Calcium imaging

Stage 6 cell clusters were stained with 20 μM Fluo-4 AM (ThermoFisher; F14201) in 2 mM glucose in KRB for 45 min at 37 °C, washed repeatedly, and loaded into the microfluidic device for analysis. The device was set up on a Zeiss Axio Zoom V16 Macroscope with a heated stage and fluorescence images taken with a Hamamatsu CCD camera at 50X magnification. KRB buffer was flowed at 0.5 mL/min at 2 or 20 mM glucose or at 20 mM plus 30 mM KCl. The time series images were present as a stack of multiple images which were analyzed as a hyperstack using ImageJ/Fiji software. They were first corrected for translational movement using the StackReg feature, after which they were analyzed with Regions of Interest (ROI) for each entire islet cluster, and a fluorescence intensity profile was generated using the data, with background noise subtraction. Flow-thru was also collected for insulin quantification.

### Statistical analysis

GraphPad Prism was used to calculate statistical significance. Two-sided paired or unpaired *t*-tests were used. Error bars represent s.e.m.

